# Resting Cortical Alpha and Beta Oscillations are Continuous Across Space and Frequency

**DOI:** 10.64898/2026.09.22.752811

**Authors:** Isabel S. Wilson, Santiago I. Flores-Alonso, Alex I. Wiesman

**Author notes:** Corresponding author Correspondence: Alex I. Wiesman, PhD, Department of Biomedical Physiology & Kinesiology Simon Fraser University, Burnaby, British Columbia.

## Abstract

Cortical alpha and beta oscillations are traditionally conceptualized as spatially distinct, with alpha rhythms constrained to the occipital cortex and beta rhythms to frontal cortices. Little work has tested this spatial separability empirically and explored whether the distribution of these cortical rhythms may be continuous in space and frequency. Using task-free magnetoencephalography of 604 participants and density-based unsupervised clustering of individual spatio-spectral peaks, we show that the distribution of cortical oscillatory activity across the alpha-to-beta frequency range is structured along a continuous spatio-spectral gradient from occipitotemporal to frontal cortices. Across individuals, the expression of this gradient correlates positively with cognition and decays with age. This work introduces a continuous spatio-spectral organizational structure linking alpha- and beta-band activity across all lobes of the neocortex, with a potential role in flexible segregation and integration of communication between sensorimotor regions of the brain.

## INTRODUCTION

The brain’s function depends on flexible control of communication between neurons across specialized regions (*1*, *2*). One candidate mechanism for coordinating this communication is neural oscillations produced by the synchronous firing of neurons (*3*). Periodic rises and falls in these oscillations create temporal windows of high and low sensitivity to stimulation, which facilitates or restricts communication between connected populations of neurons (i.e., frequency coupling (*4*)). Neurons have a higher likelihood of synchronizing with their neighbors (*4*); importantly, they also have a higher likelihood of synchronizing with neurons exhibiting similar oscillatory frequencies (measured in cycles per second, or Hz), giving rise to frequency-specific oscillating networks across the cortex (*5*). Though most distinct during active task processing, oscillatory coordination is also exhibited during task-free or so-called “resting” states. In different cortical regions, populations of neurons oscillate at distinct frequencies, creating a topography of “characteristic frequencies” across the cortex. Resting-state oscillations thus create a framework for coordinated communication on a static anatomical scaffolding that forms the basis for integration across regions during active cognition.

The canonical view is that oscillations aggregate into discrete frequency bands which act relatively independently, and which are each associated with different anatomical generators and behavioural states. For example, oscillations at the alpha frequency (8—12 Hz) are strongest in occipital regions and are of interest to visual researchers (*6*, *7*), while oscillations at the beta frequency (15—30 Hz) are strongest in fronto-parietal regions and are associated with motor and cognitive function (*8*, *9*). Explanations of cross-regional integration, meanwhile, emphasize interactions between canonical bands, for instance through synchronization (i.e., phase coherence) within or between bands (*10–14*).

Importantly, however, it is increasingly accepted that canonical spectral boundaries are oversimplifications. Overreliance on canonical band analyses may obscure meaningful variation in the boundaries between bands across individuals, in the generating mechanism across species, and in the functions of bands across brain locations (*15*). Explorations of oscillatory patterning have also emphasized dominant oscillatory features—that is, the peak frequencies that stand out most sharply in the data—potentially causing subtle, dynamic, patterning to be ignored, analogous to how the mu rhythm was at first obscured by neighboring high-power alpha and beta (*16*), or how the default mode network was initially thought to be unitary (*17*).

Recent studies have sought to describe patterns of intrinsic oscillatory activity in a more data-driven way but have still largely constrained analyses by canonical frequency bands. For example, in a recent application of unsupervised clustering to vertex-wise oscillatory power from a large sample of task-free magnetoencephalography (MEG) recordings, the topography of intrinsic oscillatory activity exhibited apparently sharp boundaries between the spatial distributions of canonical bands (*18*). However, these boundaries were necessitated by the analytical approach: k-means clustering requires that every data point be assigned to a cluster, and so in this application, it identified boundaries that best separated the oscillatory profiles spatially, even if continuous gradients would have provided a more parsimonious explanation of the data. Indeed, recent work (*19*) described several posterior-to-anterior gradients of oscillatory organization across the brain. In a large sample, for each region of the cortex, the authors identified the peak frequency within each band and found that peak frequencies shifted systematically from occipital to frontal regions. Though constrained within canonical bands and limited to linear trajectories along semi-arbitrary anatomical coordinates, these gradients were associated with variations in cortical thickness and aligned to the cortical sensory-to-association hierarchy (*19*). A subsequent MEG study replicated this gradient (*20*), and later electroencephalography work showed that it forms and dissolves over sub-second intervals (*21*) and is involved in inter-regional synchronization (*13*).

Though continuous spatio-spectral gradients have been separately identified within canonical bands (*19*), the idea that these gradients could link oscillations at frequencies traditionally belonging to different canonical bands has, to our knowledge, not yet been explored. Further, these gradients have been defined along semi-arbitrary axes defined by anatomical coordinate spaces that may not best describe the organization of the cortex. In this study, we characterized the distribution of oscillatory peaks across the cortex using MEG, while allowing for identification of spatial gradients that span canonical frequencies. This was done using a spectrum-first approach, where we began with a full set of frequency-wise oscillatory maps (i.e., from 8—30 Hz, in 0.25-Hz increments) and identified the spatial location of peak power for each frequency. We grouped these frequency-wise spatial peaks of oscillatory activity using a density-based clustering technique that allows for linkage in both frequency and space, unconstrained by alignment to anatomical axes. In a large sample of participants, we identified a continuous gradient linking occipitotemporal to frontal cortices across the alpha-to-beta frequency range and replicated this effect in an independent cohort. This spatio-spectral gradient comprised more than 60% of all identified oscillatory peaks and was distinct from a focal occipital-alpha cluster. Individuals varied in their expression of the spatio-spectral gradient, and this variation was associated with age and cognitive function, such that weaker gradient expression was associated with older age and worse cognitive performance.

## RESULTS

### Cortical Oscillatory Activity is Organized along a Continuous Alpha-to-Beta Spatial Gradient

Consistent with an established literature, MEG source maps of aperiodic-corrected oscillatory activity group-averaged over participants and canonical frequency bands showed two spatially discrete aggregates of oscillatory activity: alpha (8—12 Hz) in bilateral occipital regions and beta (15—30 Hz) in superior fronto-parietal regions (Fig. S1; task-free MEG from N = 604 healthy participants; Cambridge Centre for Ageing and Neuroscience [Cam-CAN] (*22*)). By resolving these maps into continuous frequency estimates of aperiodic-corrected oscillatory activity, however, we instead observed a continuous spatio-spectral gradient linking these alpha and beta peaks. That is, from approximately 8 to 30 Hz, the spatial peak of oscillatory power appeared to shift from posterior to anterior cortices with increasing frequency (Video S1). This surprising observation raised the question of whether alpha and beta oscillations might be spatially contiguous rather than discrete across the cortex (Fig. 1A). To explore this idea, we represented spatio-spectral patterns of oscillatory activity as “trajectories” using the peak spatial location of aperiodic-corrected power from alpha through beta in steps of 0.25 Hz (Fig. 1B)—an approach that combines the conventional use of peak power as a representative measure of oscillatory activity with highly spatially-sampled spectral parameterizations to define the spatial focus of oscillatory activity at each frequency. We pooled these spatio-spectral trajectories across individuals in our large dataset and subjectively observed that frequency increased steeply along the posterior-to-anterior and lateral-to-medial axes, suggesting that a spatio-spectral oscillatory gradient was present at the group level (Fig. 2A).

**Fig. 1.**
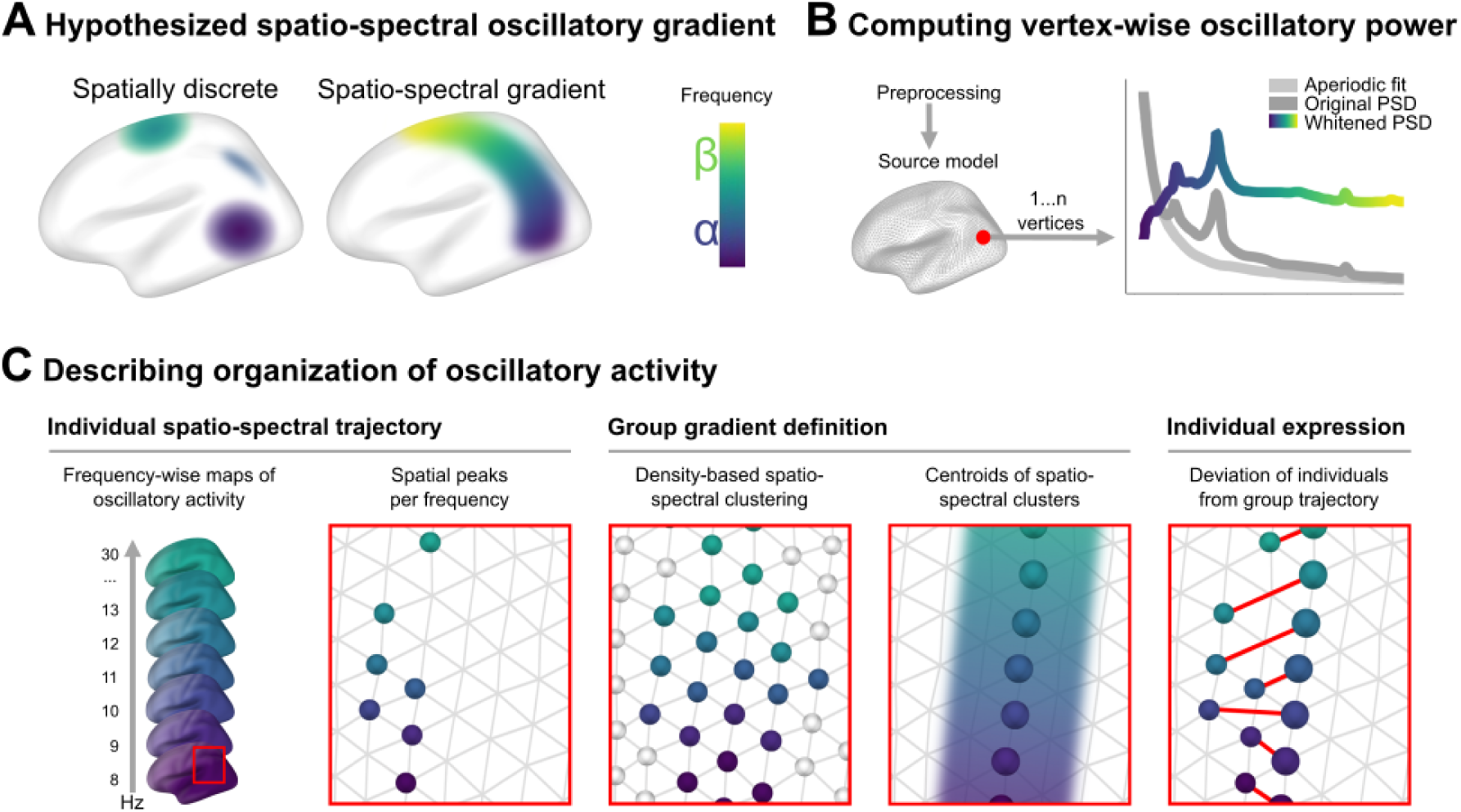
Overview of hypothesis and methods. **(A)** Conceptual illustration of study hypothesis: Oscillatory power shown as spatially discrete (traditional) versus organized along a continuous spatio-spectral gradient (hypothesized) across the cortex. **(B)** Schematic of the pipeline for computing frequency-resolved oscillatory power per participant. After standard preprocessing and source modelling, we computed the power spectral density (PSD) at each vertex, then parameterized and removed the aperiodic component. The resulting “whitened” PSD was used for further analysis. **(C)** Schematic of the pipeline for quantifying the spatio-spectral organization of oscillatory activity. From left to right: For each participant and each frequency from 8 to 30 Hz, we generated spatially resolved maps of oscillatory activity in 0.25-Hz increments. We then computed the spatial coordinates of peak oscillatory power from each of these maps. Collating these peak locations yielded a frequency-resolved trajectory of peak oscillatory power per participant. To define the spatio-spectral gradient at the group level, we pooled trajectories from a large sample of participants (N = 604), then applied density-based spatio-spectral clustering to obtain clusters of peaks linked in space and/or frequency. We summarized each cluster by computing the spatial centroid of member peaks at each frequency, thus obtaining a group gradient trajectory. To quantify the extent to which individuals “expressed” the group gradient—that is, the degree to which each individual’s trajectory of spatio-spectral peaks tracked the gradient identified at the group level—we computed the mean Euclidean distance between individual peaks and the group trajectory, rescaling these distances by subtracting them from the length of the coordinate axis.

**Fig. 2.**
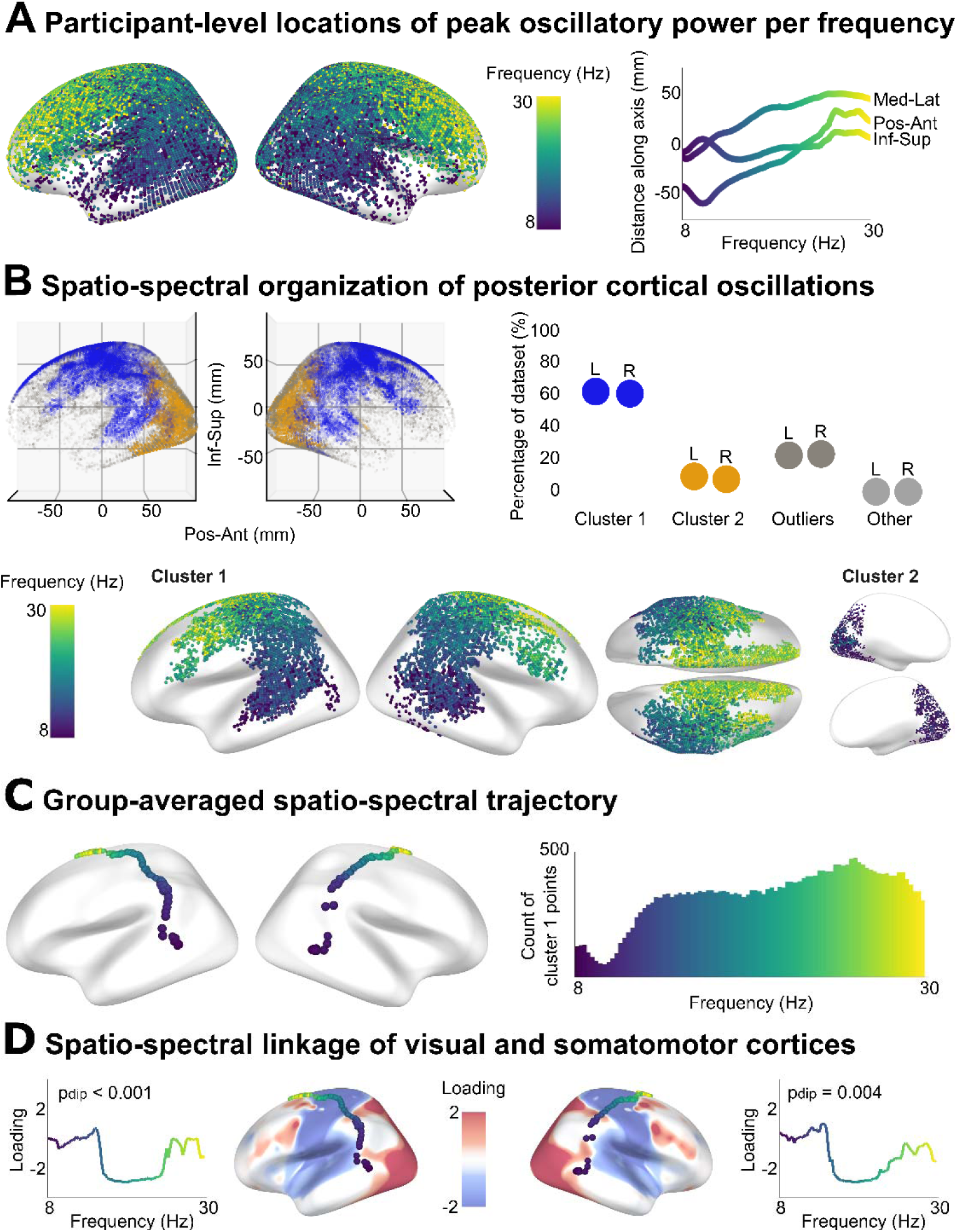
Density-based spatio-spectral clustering of cortical oscillatory activity. **(A)** In our Cam-CAN sample (N = 604), participant-level locations of peak oscillatory activity per frequency were distributed across the cortical surface, with an apparent increase in frequency observed along the posterior-anterior (Pos-Ant) axis, as well as a weaker increase along the medial-lateral (Med-Lat) axis, and no clear trend along the inferior-superior (Inf-Sup) axis. **(B)** Density-based unsupervised clustering of spatio-spectrally-linked oscillatory peaks identified four broad categories: cluster 1 (the largest cluster; colored blue), cluster 2 (the second-largest cluster; colored orange), outliers (colored dark grey), and many clusters of negligible size (179 in the left hemisphere and 211 in the right, each accounting for < 0.5% of peaks; colored light grey). Cluster 1 points were dense across the posterior cortex, showing a clear spectral trajectory from the occipitotemporal lobes (alpha frequencies) to the frontal cortices (beta frequencies). In contrast, cluster 2 points were confined to occipital regions and low frequencies. **(C)** To better describe the distribution of peaks in cluster 1, we computed the spatial center of mass of peaks per frequency, obtaining a group-averaged spatio-spectral trajectory for each hemisphere. We note that the number of points included in the cluster differed across frequencies, with a larger number of peaks included between the high-alpha and beta bands. **(D)** We mapped the group-averaged trajectory onto the second principal component of the cortical functional hierarchy, in which somatosensory/motor and auditory regions are represented by positive loadings and visual regions are represented by negative loadings (*24*). Hartigan’s dip test rejected unimodality of the distribution of loadings along the gradient trajectory, suggesting that the trajectory traverses multiple primary sensory cortices.

To quantify this observed spatio-spectral oscillatory gradient at the group-level, we adapted an established spatio-temporal density-based clustering algorithm (ST-DBSCAN (*23*); Fig. 1C) to the spatial locations of participant-wise oscillatory peaks in the frequency domain. Consistent with a continuous spatio-spectral organization, the majority of oscillatory peaks (left: 62.81%; right: 62.02%) fell into a single large cluster (cluster 1; Fig. 2B), which spectrally-linked occipitotemporal-to-frontal cortices across the alpha-to-beta frequency range (8—30 Hz; Fig. 2C). We additionally observed a smaller “alpha-occipital” cluster (cluster 2; left: 9.97%; right: 8.23%) that was mostly composed of peaks at lower frequencies (left: 80% of points were between 8 and 11.75 Hz; right: 80% of points were between 8 and 11.25 Hz) and occipital cortices, and several other clusters of negligible size (< 0.5% of oscillatory peaks; left: 179 clusters; right: 211 clusters) that were scattered across the cortex. The remainder of the points were classified as noise/outliers (left: 22.91%; right: 23.62%; Fig. 2B).

To summarize the group-level topography of cluster 1, we computed the spatial center of mass of peaks per frequency, obtaining a group-averaged spatio-spectral trajectory per hemisphere (Fig. 2C). We then contextualized this trajectory with known organizing principles of cortical function by mapping it onto the second principal component of the cortical functional hierarchy (Fig. 2D; (*24*)), which separates sensorimotor regions based on their processing modality, from visual to somatosensory/motor and auditory (*24*). The gradient trajectory was not constrained to any one sensory modality; Hartigan’s dip test (*25*) rejected unimodality (left: D = 0.13, *p*_dip_ = 0.000; right: D = 0.07, *p*_dip_ = 0.004), suggesting the trajectory traverses multiple sensorimotor cortices.

### Validation of the Spatio-Spectral Gradient in an Independent Dataset

Validation analyses used a smaller sample of task-free MEG from N = 160 healthy participants (Open MEG Archive [OMEGA] (*26*)) collected at a different site, using a different paradigm (i.e., eyes-open versus closed), and on a different instrument. From the distribution of peak oscillatory power per frequency in this validation dataset, we again subjectively observed a steep increase in frequency along the posterior-to-anterior axis (Fig. 3A). We sought to quantitatively test this gradient, but the smaller sample size of the OMEGA dataset limited cross-sample comparison of ST-DBSCAN clusters. Instead, we quantified the alignment of oscillatory peaks in the OMEGA validation sample to the spatio-spectral gradient identified in the Cam-CAN data, with an above-chance alignment reflecting external validation of the cortical spatio-spectral gradient. Alignment of OMEGA oscillatory peaks to the Cam-CAN spatio-spectral gradient significantly exceeded that observed in frequency-shuffled null distributions (*p* < 0.001; Fig. 3B). This result held across increasing bin sizes used for frequency shuffling, indicating that it is not an artifact of spatio-spectral autocorrelation (Table S1). We also repeated this procedure using oscillatory peaks derived from a shortened 15-second version of the OMEGA runs (N = 146). We again observed significant alignment to the Cam-CAN spatio-spectral gradient (Table S1), confirming that the gradient exists at relatively short timescales and is robust to effects of data duration (Fig. 3).

**Fig. 3.**
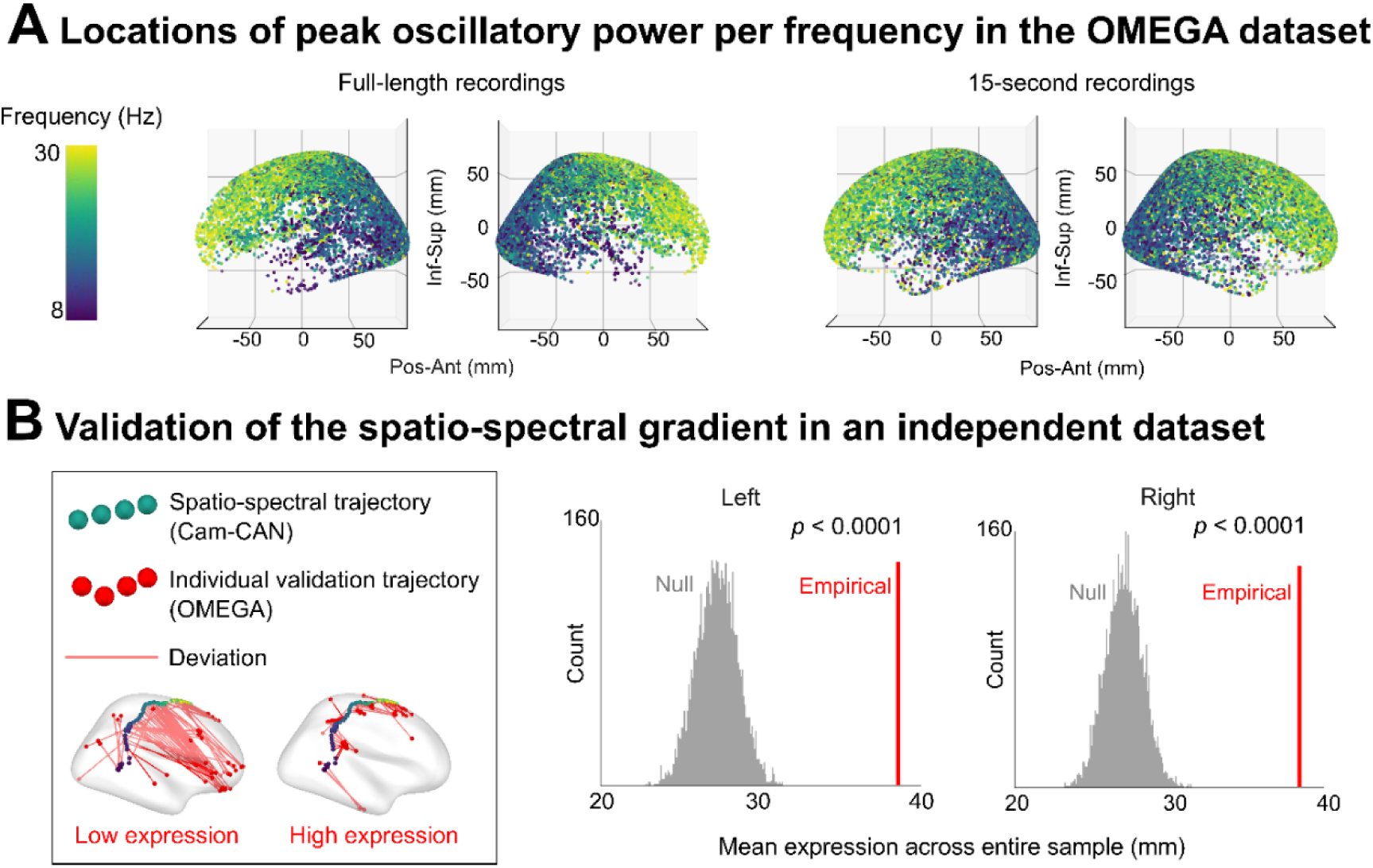
Validation in an independent dataset. **(A)** Locations of peak oscillatory power per frequency for full-length (N = 160) and 15-second (N = 146) recordings from the OMEGA validation sample. **(B)** Left: Schematic of the method used for computing alignment of individual peaks from OMEGA to the Cam-CAN spatio-spectral gradient. The Euclidean distance between the Cam-CAN group spatio-spectral gradient trajectory (green points) and the individual trajectory for each OMEGA participant (red points) is computed, resulting in a deviation per frequency per participant, which is then averaged and subtracted from a theoretical maximal deviation to generate a participant-level metric of “gradient expression.” Participants with low gradient expression have scattered frequency-wise peaks, while participants with high gradient expression have peaks distributed along a recognizable spatio-spectral trajectory. The mean of these gradient-expression values across the OMEGA sample is computed, leading to a measure of mean gradient expression. Right: Comparison between empirical OMEGA group expression of the Cam-CAN gradient (red) and a null distribution of frequency-shuffled expression values (grey).

### Expression of the Spatio-Spectral Gradient Decreases with Age and is Associated with Cognition

We tested whether individual differences in spatio-spectral gradient expression, quantified as the degree to which the distribution of individual spatio-spectral peaks tracked the group gradient trajectory (Fig. 3B), were related to age and cognitive ability. We found that gradient expression decreased with age (b = −0.053, *p* < 0.0001; Fig. 4A), and that cognitive ability increased with gradient expression beyond the effects of age (b = 0.157, *p* < 0.01; Fig. 4B). To ensure this relationship was not non-specifically associated with age-related differences in data quality, movement, or other confounds, we also tested the association between age and the overall variability in the distribution of cluster non-specific individual spatio-spectral peaks. We found no significant relationship (b = 0.026, *p* = 0.242), which confirmed that our gradient-age results are not simply due to increased spatial variability of oscillatory activity in older adults.

**Fig. 4.**
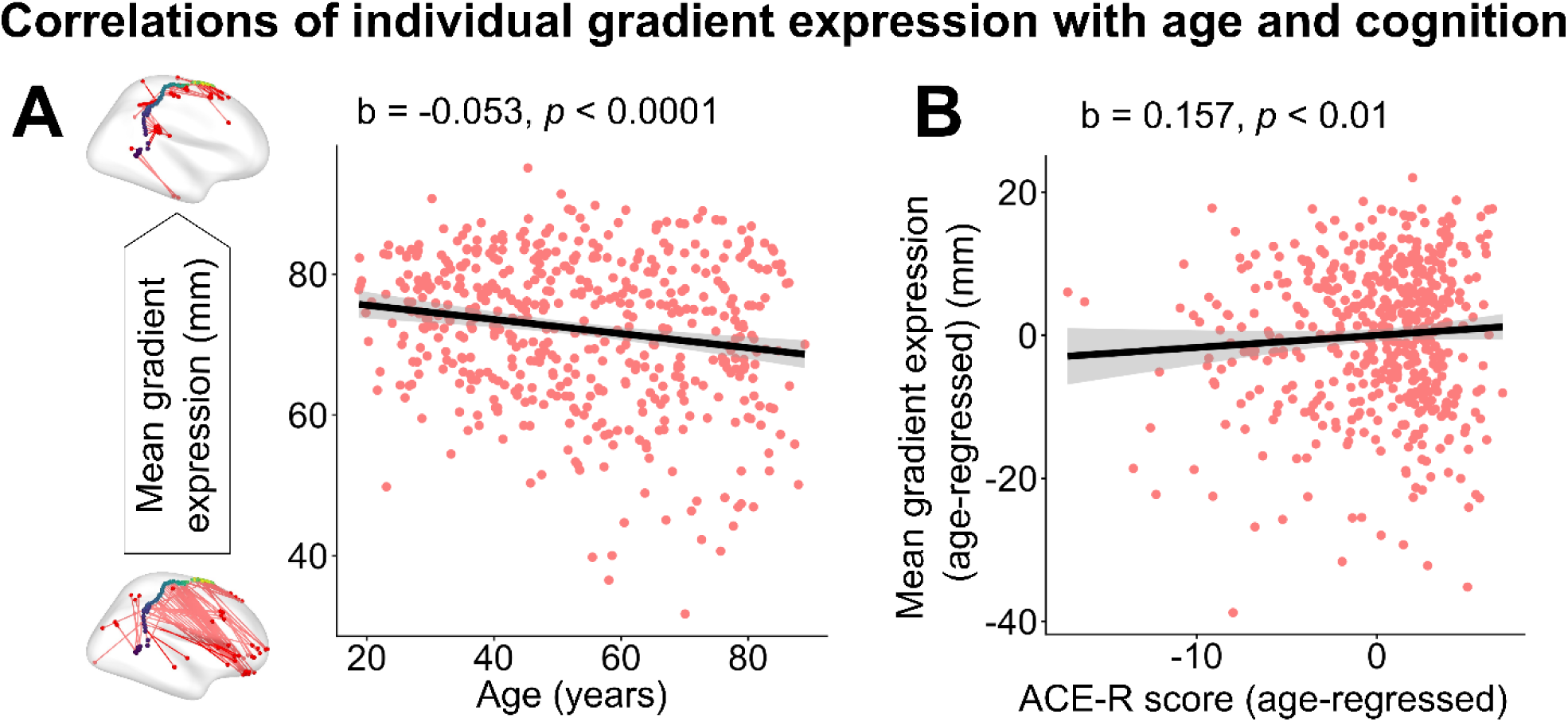
Associations between spatio-spectral gradient expression, age, and cognitive ability. Linear relationships between **(A)** mean gradient expression and age and **(B)** age-regressed mean gradient expression and age-regressed ACE-R score in the Cam-CAN dataset (N = 593). Each dot represents an individual. Grey bands represent the 95% confidence interval for the linear model fit.

## DISCUSSION

Using unsupervised density-based clustering on a large sample of source-imaged MEG data, we sought to describe the linked spatio-spectral organization of intrinsic oscillatory activity in the human cortex. Contrary to conventional knowledge that alpha and beta oscillatory activity are spatially segregated to occipital and frontal cortices, respectively (*6*, *8*), we identified a continuous spatio-spectral gradient linking occipitotemporal alpha and frontal beta oscillatory activity. Expression of this gradient varied across individuals, and this variation was associated with age and cognitive function. This gradient also replicated in an independent dataset, suggesting that it is not an artifact of site, task-free recording paradigm, or instrumentation. To our knowledge, this work is the first to introduce a continuous organizational structure linking alpha- and beta-band activity across all lobes of the neocortex.

The idea that a gradient over space and frequency could serve as an organizational structure for cortical neurophysiology has previously been explored in narrowband frequency ranges (*13*, *19–21*) and is intuitive, given that a breadth of cortical features vary along smooth continuums of the cortex, e.g., myelination, synaptic connectivity, gene transcription, and cortical thickness (*27–29*). The directionality of this gradient also generally matches previous reports of posterior-anterior structural and functional cortical organization (*19*, *27–29*), with implications for cognitive function (*13*, *30*, *31*). However, we emphasize that the spatio-spectral gradient identified in our study was not defined using semi-arbitrary anatomical axes and followed a non-linear trajectory across the cortex as frequency increased: first running inferior-to-superior from middle temporal regions to inferior parietal cortices, before orienting posterior-to-anterior and extending into prefrontal areas. This both indicates the importance of considering cortical gradients that track non-linearly across anatomical coordinate systems and highlights the utility of unsupervised density-based clustering for this purpose.

We speculate that the spatio-spectral gradient could represent a mechanism by which occipitotemporal and frontal cortices flexibly integrate information in intermediate regions, while maintaining relative functional independence between primary sensorimotor processes. Information transfer between adjacent neuronal groups is thought to rely on cross-frequency inter-regional coupling at integer phase lags (*32*). Alignment between the excitatory windows of neuronal groups enables flexible information transfer between them: when two groups’ millisecond-scale windows of excitability overlap, a spike sent by one group is more likely to arrive during a receptive phase in the other and elicit a postsynaptic potential (*14*, *32*). As such, neighboring locations on the spatio-spectral gradient would be expected to communicate more readily via the uniform short-range connectivity previously described across these regions (*33*). In contrast, more distant areas (e.g., occipitotemporal and frontal regions) would be inherently functionally segregated by their distinct spectral signatures, with information transfer relying on signal transmission along long white matter tracts (*34*), with slower conductance delays limiting flexibility. Future studies could explore this hypothesis by examining which aspects of visuomotor separation versus integration are supported by long-versus short-distance functional connectivity during a task. Non-invasive neurostimulation could also be applied to enhance and/or disrupt frequency-defined signaling along the gradient, to test whether visuomotor integration can be causally manipulated. We additionally note that the alpha spectral peaks at one extreme of this gradient were localized to the occipitotemporal cortex, with a second, independent, cluster of low-alpha peaks localized to the ventral occipital cortex. Potentially, spatio-spectral separation between primary visual and secondary occipitotemporal cortices could allow for functional separation between activity related, respectively, to basic feature encoding and higher-level processing of coherent objects and motion (*35*), with higher-order representations then being selectively linked to somatomotor cortex via the observed spatio-spectral gradient.

Gradient expression varied across individuals, consistent with well-documented individual variability in oscillatory activity (*36*). Across individuals, stronger gradient expression correlated with younger age and better cognitive performance. This result suggests that the gradient may serve a functional role in cognition, potentially via the mechanisms speculated above. However, we note that the ACE-R battery used to measure cognitive abilities was designed for dementia screening and may not represent the most sensitive test for quantifying subtle, age-related changes in cognition. Detailed characterization of individual trajectories has also been left for future work because the unsupervised clustering method used to define the gradient requires a large number of datapoints, and so is inappropriate for individual data. To describe individual-level trajectories, we instead relied on comparisons to the group average. Despite correlating with lower age and higher age-corrected cognitive function, individual-level gradient expression was essentially a measure of “averageness,” and it is possible that different patterns or “subtypes” of trajectory expression may confer different cognitive advantages. Future work could, for example, use denser sampling of individual participants, perhaps with temporally windowed data, to characterize individual trajectories while relying less on the group mean for gradient definition.

Prior work has likely failed to detect this gradient due to four limitations. First, conventional analytical approaches assume that oscillatory rhythms should be spatially separable (*18*, *37*, *38*). This is, at least partially, due to early methodological constraints on the field: the earliest neurophysiological recordings of the cerebral cortex were densely irregular single-channel time-series, necessitating the identification of unambiguous data regularities. What stood out in the time series were 10 Hz oscillations, and these oscillations were subsequently localized to the occipital cortex (*39*), leading to a trend of data analysis focusing on spatio-spectral independence. Second, even those studies that have removed the constraint of spatial separability have remained focused on canonical frequency bands. For instance, when a recent analysis decomposed 1–40 Hz oscillatory activity into a distribution of spectral peaks, the gradient reported was restricted to the range with highest power (i.e., the 7–12 Hz alpha frequency range) (*19*) even though an unrestricted analysis might have revealed more parsimonious gradients spanning other frequency bands as well. Third, until recently, separating oscillatory activity from aperiodic components has been computationally expensive, but recent advances in this area now allow for parameterizing neural spectra that are densely sampled across the neocortex (*15*). Finally, and as discussed above, the gradient we identify does not linearly track any one anatomical axis. Previous approaches have constrained spectral gradient identification to alignment with these semi-arbitrary coordinate spaces, which would obscure the rich non-linear spatio-spectral trajectory that we observe here.

Using task-free MEG, we showed that the distribution of oscillatory activity across the alpha-to-beta frequency range is structured along a continuous spatio-spectral gradient from occipitotemporal to frontal cortices. Across individuals, the expression of this gradient varies with age and cognitive function. This introduces a potential mechanism for flexible coordination of communication between regions of the brain, and we envision it as a methodological and conceptual blueprint for future studies of human neurophysiology. Our findings open exciting opportunities for new research, including those that we have already described involving task-based neuroimaging, frequency-targeted non-invasive neurostimulation, and gradient subtyping. Future work should also explore the gradient’s temporal dynamics: does this effect, like previously described frequency-limited oscillatory gradients (*13*, *21*), form and dissolve over time? And if so, how does this instability relate to its role in cognitive function? It might also be expected that this spatio-spectral organization would be re-organized or degraded in neurological disorders that affect visuo-motor integration, for example in people living with Parkinson’s disease (*40–42*).

## MATERIALS AND METHODS

### Participants and Data

Our data consisted of two samples of task-free MEG from healthy adults: a discovery sample (Cambridge Centre for Ageing and Neuroscience dataset [Cam-CAN] (*22*, *43*)) and a validation sample (Open MEG Archive [OMEGA] (*26*)). Differences in the parameters of these datasets (Cam-CAN: 204-planar gradiometer & 102-magnetometer MEGIN/Elekta VectorView, Helsinki, Finland; 1000-Hz sampling rate; eyes-closed resting-state; OMEGA: 275-axial gradiometer CTF, Port Coquitlam, BC, Canada; 2400-Hz sampling rate; eyes-open resting-state) allowed us to test generalizability. T1-weighted structural MRIs allowed for source localization to individual anatomy. The data collection and management protocols for the Cam-CAN and OMEGA repositories were approved by the research ethics boards at the University of Cambridge (reference: 10/H0308/50 (*22*)) and Montreal Neurological Institute (*26*), respectively, and secondary analysis was approved by the research ethics board at Simon Fraser University (reference: 30002675). All participants in both studies provided written informed consent in accordance with the Declaration of Helsinki.

From Cam-CAN, we excluded individuals for whom MEG or structural MRI data failed preprocessing or source reconstruction quality control. This left 604 task-free acquisitions (593 had demographic data; for these, age range = [18-90]; mean age = 55.30 ± 18.07 years; 302 males), each approximately 8 min. From OMEGA, we selected participants with no noted neuropsychiatric or neurological diagnoses and with complete MEG coregistration data, for a sample of 192 healthy individuals. The sample was further constrained to exclude short (< 5 min) task-free recordings and data that failed MEG preprocessing or source reconstruction quality control—leaving a final sample of 160 participants (mean age = 29.97 [SD = 10.91]; age range = 18.19-74.68 years; 104 males; task-free recording length range 5-10 min, mean 6.05 min, SD 2.00 min).

### MEG Data Processing

MEG data were preprocessed in MNE-Python 1.10.2 ((*44*); Python 3.13.9) following good-practice guidelines (*45*). For Cam-CAN data, spatiotemporal signal-space separation (tSSS; (*46*), with window duration = 10 s, correlation limit = 99%) was applied to task-free data. Both datasets were band-pass filtered (Cam-CAN: 1–90 Hz; OMEGA: 0.5-200 Hz), notch-filtered (Cam-CAN: 50 Hz; OMEGA: 60, 120, 180 Hz), downsampled (Cam-CAN: 500 Hz; OMEGA: 600 Hz), and cropped to a fixed interval across participants (Cam-CAN: 300 seconds; OMEGA: 240 seconds). Note that these crop lengths exceed minimum data-duration recommendations for stable parameterized features in these datasets (*47*). We used electro-cardiogram and -oculogram recordings to define signal-space projections (*48*) for the removal of cardiac and ocular artifacts. For both datasets, bad segments were identified on the basis of deviation from the median signal amplitude and deviation from the numerical gradient of signal amplitude (over 1-second windows), but not considered in computation of the data covariance for source imaging. Spatial filters were normalized using a unit-noise-gain constraint to reduce depth bias and ensure comparable sensitivity across cortical locations. For both datasets, empty-room recordings acquired close in time to each participant’s task-free scan and lasting at least two minutes were processed using the same pipeline (excluding tSSS in the Cam-CAN dataset) and used for noise statistics in source analysis.

MEG data were coregistered to individual segmented T1-weighted MRIs (*FreeSurfer 7.4.1 recon-all* (*49*)) using publicly available transformation matrices (Cam-CAN: (*50*); https://github.com/tbardouille/camcan_coreg; OMEGA: https://www.mcgill.ca/bic/neuroinformatics/omega), which had been precomputed using fiducial landmarks (nasion and bilateral preauricular points) and digitized head surface points. The process was the same for both datasets: forward solutions were computed using a three-layer Boundary Element Method (BEM) head model (*51*). Cortical source activity was reconstructed using linearly constrained minimum-variance (LCMV) beamforming (*52*). Data covariance matrices were estimated from the preprocessed sensor-level data and regularized (λ = 5%) to stabilize spatial filter estimation. Beamformer weights were computed across 20,484 cortical vertices per participant, with source orientations optimized in the direction of maximal projected power.

### Computation of Individual-level Spatial Oscillatory Peaks per Frequency

Per participant, we computed the spatial location of peak oscillatory (i.e., aperiodic-corrected) spectral power at each frequency from 8 to 30 Hz in 0.25-Hz increments. To do so, we first obtained vertex-wise estimates of the power spectral density (PSD) using Welch’s method (4-s time window and 50% overlap for both datasets (*53*)). Next, we subtracted the aperiodic component of oscillatory activity from the PSD. The aperiodic component had been computed vertex-wise using the specparam/FOOOF algorithm (*54*) implemented in Python (frequency range = 1-40 Hz; peak width limits = 0.5-12 Hz; maximum n_peaks_ = inf; minimum peak height = 0 dB; proximity threshold = 2 SD; fixed aperiodic). Note that gamma frequencies (> 40 Hz) were not included in this analysis, as the FOOOF algorithm struggles with fitting PSD data properly above 40 Hz due to the loss of linearity in log-log space and lack of clear peaks in gamma frequency ranges and above (*54*).

Once we had spatial maps of oscillatory power at each frequency bin, we computed the spatial coordinates of the peak power per bin using the *get_peak()* function in MNE-Python (*44*). This resulted in 88 peaks per participant per hemisphere, one for each 0.25-Hz increment from 8 to 30, distributed along the cortical surface. In total there were 88×604 such peaks per hemisphere in the Cam-CAN dataset and 88×160 per hemisphere in the OMEGA dataset.

### Spatio-Spectral Density-Based Spatial Clustering of Oscillatory Peaks

To determine whether these frequency-wise oscillatory peaks were organized along a spatio-spectral cortical gradient in the Cam-CAN dataset, we performed clustering with Spatio-Temporal Density-Based Spatial Clustering of Applications with Noise (ST-DBSCAN). This extension of the widely used DBSCAN algorithm incorporates an additional user-defined dimension (*23*) for detecting clusters, typically time, which we substituted here for frequency. ST-DBSCAN assigns points to clusters based on local density, allowing for the identification of clusters with irregular shapes and the detection of outliers. A point is assigned to a cluster if it satisfies predefined density criteria determined by three user-defined parameters: the minimum number of neighboring points required to form a cluster (*minPts*), the maximum spatial distance between neighboring points (*eps1*), and the maximum frequency distance between neighboring points (*eps2*).

In our application of ST-DBSCAN, we selected initial *minPts* and *eps1* parameters following heuristic recommendations (*23*), then optimized these parameters using a grid search. The heuristic involves first defining *minPts*, then using *minPts* to find *eps1*, with *eps2* selected based on the resolution of the temporal/spectral data. As recommended, we first defined *minPts* as the natural logarithm of the total database size (*minPts* = 10). Then, we determined the spatial Euclidean distances to the *k*-nearest neighbours (*k* = minPts) for each centroid, plotted these distance values in descending order, and set *eps1* to approximately the distance defined at the elbow of the plot (*eps1* = 0.03125). Based on the spectral resolution of our data, we selected a starting value for *eps2* of 0.5. We ran a grid search around these starting values; that is, for each possible combination of values, we ran ST-DBSCAN and computed the Davies-Bouldin index (*55*) for the resulting clusters. We selected the parameters that minimized the Davies-Bouldin index (*minPts* = 9, *eps1* = 0.0319, and *eps2* = 0.5). Having selected these parameters based on the left hemisphere data, we used them to run ST-DBSCAN on the data from both the left and right hemispheres. For both hemispheres, the frequency-resolved centroids of the peaks belonging to a cluster were used to represent a group summary measure of that cluster’s spatio-spectral gradient trajectory. We contextualized the spatio-spectral gradient trajectory of cluster 1 by extracting the value of the loading of principal component 2 of the cortical functional hierarchy (*24*) at each frequency-linked centroid location, and testing for unimodality of the resulting frequency-wise trajectory of loadings using Hartigan’s dip statistic (*25*).

### Replication of the Spatio-Spectral Gradient in an Independent Dataset

Having identified an occipitotemporal-to-frontal spatio-spectral gradient in the Cam-CAN dataset, we next sought to validate this gradient in the OMEGA sample. The smaller size of our OMEGA sample would preclude the use of ST-DBSCAN with comparable clustering parameters, and so instead we quantified the degree to which the distribution of individual peaks from the OMEGA validation sample tracked the spatio-spectral gradient trajectory identified in Cam-CAN. That is, from 8 to 30 Hz, we computed the Euclidean distance of each OMEGA participant’s frequency-wise individual peaks from the corresponding centroids of the Cam-CAN trajectory; we then subtracted these distances from a theoretical maximum value (i.e., the length of the FreeSurfer coordinate axis—100 mm), and the mean of these differences represented overall alignment to the gradient. We compared these mean-gradient-expression values to null distributions constructed from mean-gradient-expression values computed on frequency-shuffled data, with a significance threshold of *p*PERM < 0.05. To account for spatio-spectral autocorrelation, we recomputed these statistics with frequency-shuffled data with increasing bin sizes (bin sizes: 0.25 Hz, 0.5 Hz, 1 Hz, and 2.75 Hz).

We also sought to test the stability of the gradient over shorter data segments. We created shortened task-free runs by extracting the first continuous 15-second segment of clean data from each of our OMEGA participants. These segments were available for 146 participants. We repeated the mean-gradient-expression analysis described above for these shortened segments.

### Associations with Age and Cognition

Finally, we asked whether individual differences in expression of the spatio-spectral gradient were associated with individual differences in age and cognitive ability. For this, we used the Cam-CAN dataset because it includes an even distribution of ages and detailed neuropsychological testing data from Addenbrooke’s Cognitive Examination (ACE-R (*56*)). The ACE-R exam comprises attention/orientation, memory, verbal fluency, language, and visuospatial domains, with total score used here as a measure of domain-general cognitive ability (*56*). For the 593 participants with complete age and ACE-R data, we regressed mean-gradient-expression values (that is, inverted Euclidean distance from each participant’s cluster-1 peaks to the group cluster-1 centroid) against age to examine age-related changes in expression of the spatio-spectral gradient. We then regressed age out of total ACE-R scores and mean gradient expression to obtain age-independent measures of individual-level cognitive abilities and gradient expression, respectively, and tested for associations using linear regression of these two variables. As a control, we created an analogous metric of cluster non-specific individual variability by computing the Euclidean distance from individuals’ frequency-wise peaks to the corresponding group centroids for each respective frequency, then subtracted these distances from 100, and averaged them within individual. We regressed this metric against age to determine whether our cluster-specific age effects were attributable to age-related confounds such as data quality and movement artifacts.

## Author contributions

Conceptualization: A.I.W.

Methodology: A.I.W., S.I.F.

Investigation: I.S.W.

Visualization: I.S.W., S.I.F.

Supervision: A.I.W.

Writing—original draft: I.S.W.

Writing—review & editing: A.I.W., S.I.F.

## Data, code, and materials availability

All data needed to evaluate the conclusions in the paper are present in the paper and/or the Supplementary Materials. Data used in this work were obtained from the Cam-CAN repository (available at http://www.mrc-cbu.cam.ac.uk/datasets/camcan/ (*22*, *43*)) and the OMEGA repository (available at https://www.mcgill.ca/bic/resources/omega (*26*)). Code used to perform the analysis is available on GitHub (https://github.com/isabel-wilson/SpatioSpec_code).

## SUPPLEMENTARY TEXT

**Fig. S1.**
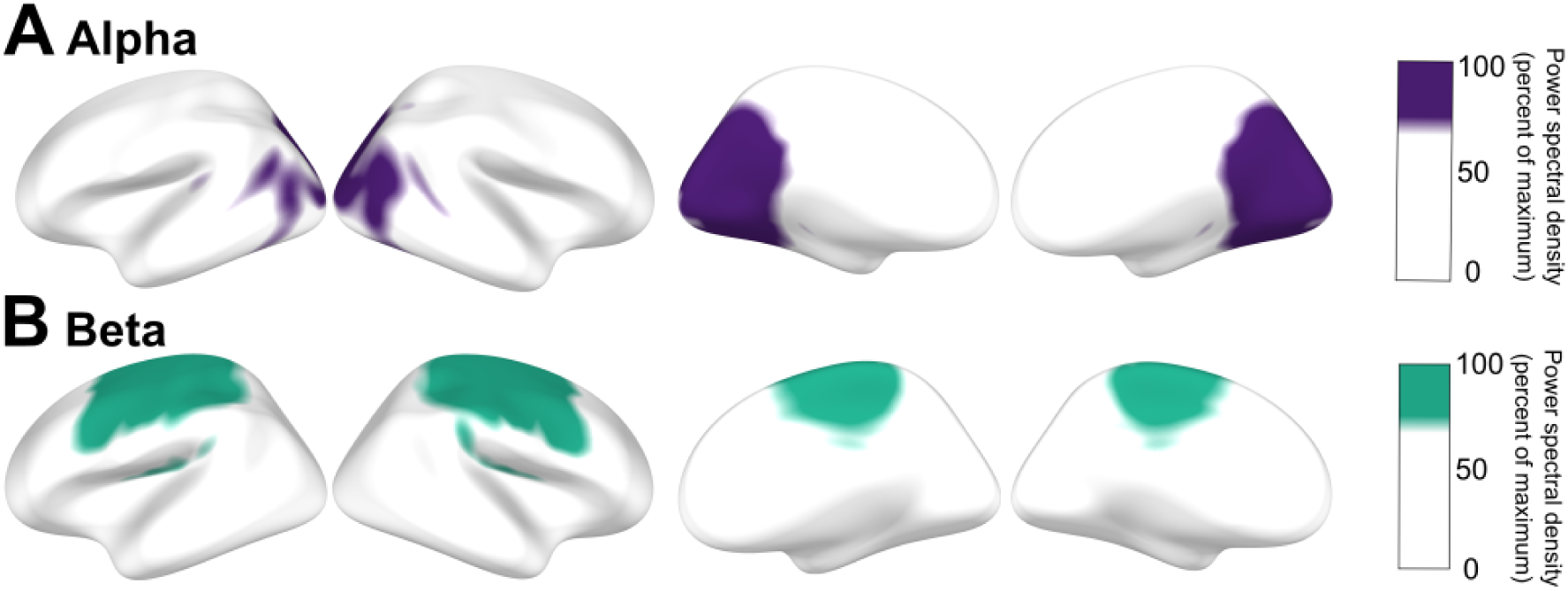
Distribution of alpha and beta oscillatory activity. MEG source maps of aperiodic-corrected oscillatory power averaged over N = 604 participants (Cambridge Centre for Ageing and Neuroscience [Cam-CAN] (*22*)) and the canonical frequency bands alpha (8—12 Hz) and beta (15—30 Hz).

**Table S1.** Empirical versus frequency-shuffled mean gradient alignment.

|  | **Full sample** |  |  |  | **First 15s** |  |  |  |
|---|---|---|---|---|---|---|---|---|
|  | **Left** |  | **Right** |  | **Left** |  | **Right** |  |
| **Empirical** |  |  |  |  |  |  |  |  |
|  | 38.45 |  | 37.41 |  | 31.26 |  | 30.27 |  |
| **Frequency-shuffled null** |  |  |  |  |  |  |  |  |
| **Bin size (Hz)** | **[min, max]** | ***p*** | **[min, max]** | ***p*** | **[min, max]** | ***p*** | **[min, max]** | ***p*** |
| 0.25 | [22.82, 31.41] | *p* < 0.0001 | [23.22, 31.47] | *p* < 0.0001 | [17.64, 26.14] | *p* < 0.0001 | [18.08, 25.37] | *p* < 0.0001 |
| 0.5 | [20.71, 33.68] | *p* < 0.0001 | [20.97, 33.77] | *p* < 0.0001 | [15.81, 26.82] | *p* < 0.0001 | [16.65, 26.57] | *p* < 0.0001 |
| 1.0 | [18.43, 35.26] | *p* < 0.0001 | [18.77, 35.27] | *p* < 0.0001 | [13.88, 29.58] | *p* < 0.0001 | [14.57, 28.51] | *p* < 0.0001 |
| 2.75 | [16.74, 38.97] | *p* = 0.0014 | [17.42, 37.74] | *p* = 0.001 | [12.35, 31.31] | *p* = 0.0002 | [12.86, 30.86] | *p* = 0.0012 |

Units in x-y-z FreeSurfer coordinates.

**Movie S1.** Spatial distribution of frequency-resolved oscillatory power from 8—30 Hz. Each frame *f* shows the top 20% of whitened oscillatory power at frequency *f* in the Cam-CAN (N = 604) group-averaged source map. The blue centroid shows the center of mass of this thresholded activity.

